# Loss of precise auditory sampling as a sign of value-driven visual attentional capture

**DOI:** 10.1101/2024.04.07.588471

**Authors:** Rodrigo Caramés Harcevnicow, Thaiz Sánchez-Costa, Alejandra Carboni, Francisco Cervantes Constantino

## Abstract

Conditioning sensory signals endows persistent salience that influences attention, even after they no longer connect with rewards. As distractors shape multisensory target processing, conditioning phenomena remain poorly understood in relation to continuous sensory encoding. We investigate the effects of value-driven attentional capture by visual cues on the ability to reliably phase-lock cortical activity to temporal modulations of sound in audiovisual (AV) displays. Listening to periodically-modulated sound, observers discriminated between two visual object streams flickering at different rates, searching for an AV match. Also in view were peripheral color cues, used to evaluate how they modulate participant tracking fidelity as a result of color-reward associative training. Behavioral and electroencephalography (EEG) recordings (*N*=31) show that performance impoverished in presence of colors previously associated with reward, and so did tracking reliability measured by phase locking of AV responses. Decreased temporal precision predicted participants’ reward-driven distraction, evidencing the attentional shift away from the multimodal target timing structure. Loss of consistency was furthermore present in auditory response estimates, suggesting value-driven attentional capture withdraws cortical tracking fidelity across the senses. The findings are consistent with inter-modal competition at times when incentive salience cues overtake top-down tracking of multisensory streams connected in time.

## 1. Introduction

The precision with which the brain samples a scene across time has topical relevance when sensory systems are under limiting pressure, such as in age-related sensory loss. Still, other less demanding but more pervasive lifetime conditions modulate the precision with which our multisensory scenes are resolved over time. Value-driven attentional capture (VDAC) for instance encapsulates the experience where rewarded aspects of our recent past involuntarily influence selection of sensory information (***Anderson, 2019***; ***Anderson et al., 2011***, ***2021***; ***Theeuwes & Belopolsky, 2012***; ***Belopolsky, 2015***; ***Theeuwes, 2018***). Because the old tangible benefits no longer help updated goals, the phenomenon illustrates the constraint that contextual backgrounds place upon how we sample the sensory environment ( ***Todd & Manaligod, 2018***). Specifically, VDAC adjusts the share of transient neural processing dedicated to current targets to favor ‘has-been’ cue types (***Rusz et al., 2020***).

A limited number of studies has specifically investigated how VDAC penetrates through different sensory modalities than current demands require. Using functional magnetic resonance imaging (fMRI), one of the few that did demonstrated the reshaping of visual cortex representations in presence of sounds with a rewarded history (***Pooresmaeili et al., 2014***). Following their brief stimulation design, these findings left open the question of how multimodal streams that unfold over time through common temporal dynamics could remain robust or alternative molded further by reward salience (***Hickey et al., 2010***). Notably, time features are central to natural associations promoting binding of multisensory streams into stable objects (***Bizley et al., 2016***).

Does capture by incentive salience shape the ability to engage with important elements of audiovisual scenes across time? In normal conditions, the brain entrains precisely its cortical network activity to multiple component features corresponding to different visual and auditory dynamic streams, and builds multimodal associations. A prime form of entrainment of cortical populations takes place in specialized auditory primary and secondary areas following slow (<12 Hz) amplitude modulations of sound. With a relatively high temporal precision, auditory cortical entrainment becomes yet molded by the dynamics of voluntarily attended visuals that are temporally coherent, turning them to greater multimodal consistency, as shown by both behavioral and electrophysiological studies (***Atilgan et al., 2018***; ***Maddox et al., 2015***). To date, it remains unclear if bottom-up influences similarly modulate entrainment or whether they are prevented, e.g., via reactive suppression mechanisms (***Klink et al., 2023***). Hinting to both uni- and cross-modal interactions, fMRI data shows that the same region relaying cross- and supramodal input to auditory cortex, the temporo-parieto-occipital junction (***Rolls et al., 2022***), may be directly involved in audiovisual (AV) manifestations of VDAC (***Pooresmaeili et al., 2014***; ***Antono et al., 2023***).

Attentional capture is theoretically framed as the outcome of the competition between distinct goal, saliency, and prior selection biases, each related to distinguishable mechanisms (***Theeuwes, 2025***; ***Theeuwes, 2018***; ***Belopolsky, 2015***). Biases that influence early preattentive processing such as reward history typically curb the ability to engage attentional control; by contrast, top-down biasing a specific location facilitates suppression of salient distractors elsewhere (***Theeuwes, 2025***). Importantly, the explanatory power of these models depends on the temporal window in which competition operates, and empirical evidence has traditionally been based on transient event designs. In continuous multimodal stream processing however, ongoing top-down selection may need to promote sustained changes across the timing of cortical electrophysiology. For instance attention aligns more precisely the timing of neural activity to cross-modal input, becoming relatively less sharp and more unstable when input is unattended (***Lakatos et al., 2009***; ***Lakatos et al., 2013***).

Continuous salient visual distractors may set a recurring context unfavorable to steady tracking the temporal patterning of the scene targets. Unstable tracking may propagate cross-modally when temporal variability in phase-locked auditory neural excitability rises, such as when supramodal competition continuously favors salience cues (***Kayser, 2009***). Both in invasive and noninvasive studies, temporally precise neural locking has been operationalized using neural phase uniformity testing because it directly probes the correspondence in time of stimulus-tracking responses. Magnetoencephalography findings for instance suggest that precise auditory locking is a key determinant of perceptual performance in audiovisual contexts (***Kösem et al., 2014***). For continuous sensory processing, empirical evidence is needed on whether the flux of neural precision time-wise serves to assign streams to relevant classes, playing a computational role analog to feature-binding effects of spatial attention.

The current study aims to determine whether visual capture impacts the precision of cortical responses continuously locked to temporal modulations in AV stimuli. Specifically, we employ EEG to evaluate the impact on phase variability of audiovisual and unimodal auditory responses during AV perceptual judgments. Using an attentional task combined with a frequency tagging approach (***Nozaradan et al., 2012***; ***Baldauf & Desimone, 2014***; ***de Vries & Baldauf, 2019***; ***Tabarelli et al., 2020***), we indexed observer’s selective neural tracking precision over single-trial presentations. In a two-alternative forced choice task, subjects matched one of two central visual flicker options with a single amplitude-modulated (AM) sound. Conditioned visual distractors, shown at the periphery, conveyed past reward associations. Typically set in probabilistic designs (***Anderson et al., 2011***; ***Hickey & Peelen, 2015***; ***Pooresmaeili et al., 2014***; ***Sanz et al., 2018***; ***Cho & Cho, 2021***; ***Koenig et al., 2017***), but see (***Anderson, 2016***; ***Tankelevitch et al., 2020***), reward associative learning was addressed through probabilistic and non-probabilistic (fixed) conditioning schemes as the latter are expected to be less effective (***Rouhani et al., 2018***; ***Monosov, 2020***; ***Hart et al., 2015***). Our results provide first evidence of the ability of both visual capture setups to disrupt steady AV processing, decreasing the temporal precision of sound-locked responses throughout time.

## 2. Methods

### 2.1 Participants

Thirty-four subjects (15 female; mean age 26.4 ± 4.0 SD; 4 left-handed) with no history of neurological or psychiatric disorders volunteered in the study. All participants provided formal written informed consent. They reported normal hearing and normal or corrected to normal visual acuity. Handedness information was collected, determined by subjects’ handwriting dominance. The experiments were performed in accordance with the World Medical Association Declaration of Helsinki guidelines (see Ethics statement). For two subjects, the second half of EEG recordings were not saved in error, reducing analyses to 32 participants. Specific participant data affected by an incorrect color display (1 participant) or missing sound output feedback recordings (1 participant) were not included in analyses based on that information, as indicated in the relevant Statistical analysis sections, below.

#### 2.1.1 Ethics statement

The Ethics in Research Committee of the School of Psychology at the Universidad de la República approved the experimental procedures under ethics approval record reference 191175-000824-18 dated 7 November 2018.

### 2.2 Experimental equipment and setup

Participants were seated at 50 cm in front of a 40 cm CRT monitor (E. Systems, Inc., CA) delivering visual presentations with an 83 dpi resolution and an 85 Hz refresh rate. Audio presentations were delivered with a Sound Blaster Z sound card (Creative Labs, Singapore) in combination with a Scarlett 4i4 hub interface (Focusrite Plc, UK) and high-quality Sennheiser HD 25 headphones (Sennheiser, Germany). Neural recordings were performed using a BioSemi ActiveTwo (BioSemi, The Netherlands) 64-channel EEG system and common mode sense / driven right leg with a 10/20 layout at a 2048 Hz digitization rate. Audio signal sampling was obtained from sound interface output fed back to the ActiveTwo system through an optical link cable (ERGO Optical, BioSemi). A fifth-order cascaded integrator-comb low-pass filter with a −3 dB point at 410 Hz was applied to sensor channels online, after which all signals were decimated to 1024 Hz. Online high-pass response was fully DC-coupled. Participants adjusted the sound volume presentation to a comfortable listening level prior to starting the computerized tasks. Full experimental sessions lasted ∼2.5 h.

### 2.3 Experimental protocol overview

To generate reward associations, participants were trained on the Eriksen flanker test, a response inhibition task designed to maximize gains by suppressing objects outside of the focus of spatial attention (***Eriksen & Eriksen, 1974***). At this Training stage, target and flankers were presented along with peripheral color displays which could predict earnings via controlled color-reward mappings. Unlike visual training paradigms where success is contingent on a color search display (e.g., ***Anderson et al., 2011***), here participants discriminated within a constant, narrow spatial domain. The purpose of this was two-fold: to ensure that the target space remained comparable to that at the subsequent testing stage (see below), and that color-signaling spaces remained similar across stages.

A main AV task was subsequently used as a Testing stage to investigate subject performance and neural responses in the presence of reward-associated but irrelevant color distractors. From electroencephalography (EEG) recordings of testing stage data, modulations of temporal consistency (phase coherence) were probed via frequency tagging methods (***Grahek et al., 2021***). Prior to the main experiment, participants also performed a 2-3 minute auditory-only listening session where they were presented with long versions of the experiment sound stimuli (see Auditory stimuli, below), and instructed to listen attentively with their eyes closed. Recordings from the unimodal presentations served to inform the spatial distribution associated to auditory-evoked activity across listeners.

Main experimental sessions consisted of an initial Training task (Tr1) followed by an initial AV Testing task (Tt1), both of which represented baseline conditions without reward learning (**Figure 1A**). To address reward learning effects, participants thereafter performed a second round of Training (Tr2) and Testing (Tt2) tasks with reward feedback. Presentation and response logging were performed with PsychoPy (***Peirce, 2007***) for the Training tasks and with MATLAB R2010a (Natick, United States) for the Testing tasks.

**Figure 1.**
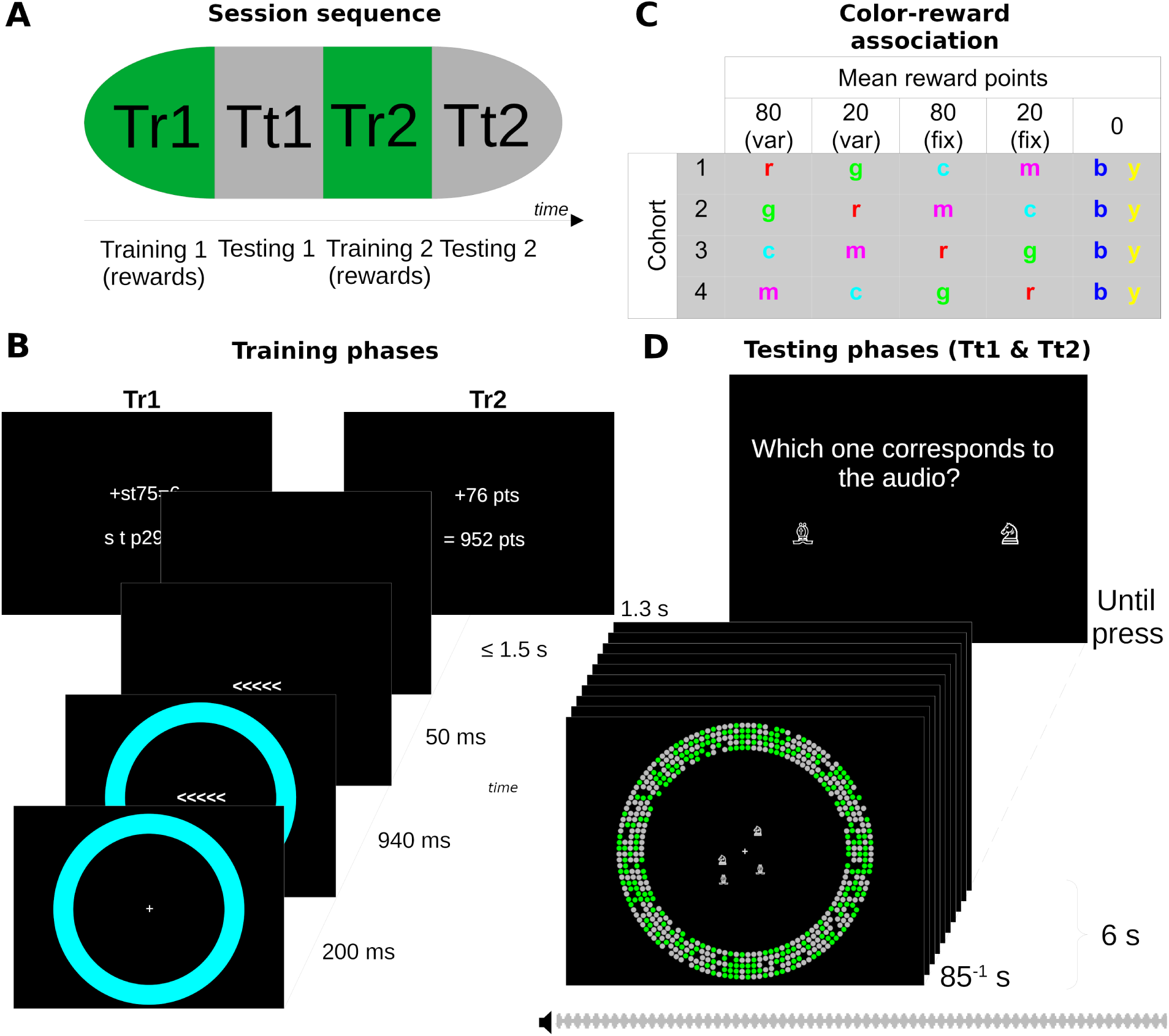
Design. **(A)** The full experiment consists of two sequential, interleaved rounds of *Training* and audiovisual (AV) *Testing* tasks designed to evaluate effects of reward learning. **(B)** At the Training stages (Tr1 and Tr2), participants accumulate points over correct trials in a standard flanker task. The colored peripheral ring may predict the range of points to be earned from correct trials. At Tr1 this information is not available (baseline) and becomes clear at Tr2 (conditioning). **(C)** Color-reward associations determine mean reward amounts and range, and are determined by cohort assigned to participants (see Materials and Methods). For instance, cohort 3 participants receive on average 80 points on correct responses if a cyan ring is also shown, and the realized amount varies across such trials. Participants do not earn points from baseline trials. **(D)** Participants are tested in a two-alternative forced choice AV task before and after conditioning, (Tt1 and Tt2, respectively), which contains a dynamic 6-second AV display consisting of an AM noise stream plus two visual symbol sets flickering at different rates. The goal is to determine which of the two sets flickers to the same sound modulation rate (9 or 11 Hz). Also included is a peripheral ringed dot pattern with flickering grey and colored dots. At Tt2, colored dots may signal reward associations learned from Tr2. See Video 1 for a full AV trial example or Supplementary Figure 1 for a frame-by-frame video sequence.

### 2.4 Training tasks (Tr1 & Tr2)

#### 2.4.1 Protocol and stimuli

Participants were asked to quickly and accurately report the direction of a central target (**Figure 1B**). Each task consisted of 216 trials with balanced congruent, incongruent, and neutral flanker conditions. A static colored ring display was shown at the start of a trial over a black background, with an internal[external] diameter subtending 21[27] degrees of visual angle (dva). Color displays were either solid red (#FF0000), green (#00FF00), blue (#0000FF), cyan (#00FFFF), magenta (#FF00FF), or yellow (#FFFF00), distributed by 36 trials per color. After an initial 200 ms of ring presentation, the task’s target and flanker symbols appeared in white, subtending 6 dva horizontally. All elements were shown for a further 940 ms, after which the ring disappeared. The symbols remained for an additional 50 ms, and there was a final 1.5 s response timeout period. Participants responded to the direction of the central symbol by pressing a keyboard button. Correct and incorrect responses were treated as valid trials whereas missed/late responses were treated as invalid.

#### 2.4.2 Reward structure

Points were awarded for valid and correct trials, by amounts determined by ring color as follows: a flat 80[20] points reward was given for valid responses in trials representing High[Low] reward and fixed conditions. In High[Low] reward and probabilistic conditions, valid responses earned a number of points that was pseudo-randomly sampled from a normal distribution (mean = 80[20] points; standard deviation = 7 points) and rounded to the nearest integer. Finally, ‘No Reward’ conditions always yielded 0 points and served as a common baseline. Colors were paired with conditions following a four cohort schedule (**Figure 1C**). Invalid (but not incorrect) trials were deducted 50 points from the accumulated score. Trial conditions and colors were presented in random order. An ad lib pause was made every 72 trials, indicating the number of accumulated points so far. The order of trials was randomized for each task. Tr1 and Tr2 were identical except on feedback presentation.

#### 2.4.3 Feedback

After entering a valid response participants received text feedback, but it was intelligible only at Tr2. For Tr1 (baseline), all characters appeared in scrambled order into uninterpretable strings. In consequence, reward-color mappings were not available for participants at the first stage. Pause-related feedback, and feedback from invalid trials which resulted in point deductions, were clearly presented in both stages. In both cases participants were instructed to accumulate as many points as possible. They were informed that the points would be converted into tokens at the end of each Testing task, to be exchanged for a prize at the end of the session. Other than through feedback, participants were not explicitly debriefed about reward-color mappings.

### 2.5 Testing tasks (Tt1 & Tt2)

In a two-alternative forced choice (2AFC) AV task, participants were presented with trials consisting of a dual visual object flicker display in tandem with a single amplitude-modulated (AM) noise sound. The goal was to correctly associate one of the two flicker object rates with the AM rate.

#### 2.5.1 Visual flicker display

The visual dual visual object flicker display consisted of a combination of two non-overlapping central (target and non-target) figure sets with peripheral distractor presentations. Central targets and nontargets. On each trial, two symbols were taken at random from a standard chess pool, i.e., king, queen, bishop, knight, rook, or pawn figures. Two replicates of each figure were shown, and flickered in phase at 9 Hz in one figure and 11 Hz in the other. Target/non-target figure designations were determined by the auditory stream (see below). The flicker rates were selected for the following technical reasons: *i*) to elicit reliable steady-state responses from visual delivery adjusted to the screen refresh rate (85 Hz) via frequency approximation (***Nakanishi et al., 2014***; Supplementary Figure 1); *ii*) for both rates to remain relatively close to the optimal low-frequency domain where auditory EEG cortical responses also lock; and *iii*) for them to be separate enough from to each other to avoid excessively difficult discrimination. The four figure locations were scattered over a circular surface extending 8 dva in diameter and shown with a steady central fixation cross (**Figure 1C**). Peripheral distractors. A peripheral ring-shaped aperture was presented with similar dimensions as in the Training task. This domain was filled with a mixture of grey and colored dots; the specific color varied per trial and was from the same color set as in Training. The ratio of grey to colored dots was fixed at 4:3 (320 grey dots), and the diameter of each dot was 0.53 dva with a density of 2.6 dots/dva^2^. Critically, to prevent the potential confound that auditory activity is differentially entrained because of mismatching dynamics between targets and distractors, peripheral dot sets flickered with colored dots exclusively synced to the target set and gray dots to the non-target set (Video 1, Supplementary Figure 1). The dots changed position every 1 s, making central figures sole reliable visual cues to perform the task. Color, figure and rates were equally distributed and balanced across trials. Dots and symbols did not overlay. Chess figures remained in their same location for 1 s before changing to different locations within the delimited central area. This resulted in a highly challenging task and then for half of the participants chess figures remained at the same initial positions throughout the trial, which was accounted for as a nuisance variable (see Data and statistical analysis). Trial stimuli displays lasted 6 seconds in total.

#### 2.5.2 Auditory stream

Two 6 s AM white noise stimuli were synthesized with 9 and 11 Hz sinusoidal modulation rates, respectively, at a 75% modulation depth and sampled at 44.1 KHz. For listening-only pre-experimental sessions (see Experimental protocol, above), two additional stimuli were created with the same conditions except that they were 54 s in duration. The same auditory stimuli were used across trials and participants.

#### 2.5.3 Audiovisual presentation

Each AV trial presentation consisted of the visual flicker display described above plus either the 9 or 11 Hz AM noise sound, and lasted 6 s. The AM rate presented at a trial defined which of the two figure flickers was the target. The onset of the noise relative to the first video frame included a uniformly distributed delay in the (0,1/ *f*) s range, where *f* is the sound AM rate, and a random phase was sampled at every trial. This manipulation allowed to discard visually-locked contributions from phase-locked activity at the inter-trial level.

#### 2.5.4 AV task and feedback

Participants were asked to detect in each trial the chess symbol whose flicker matched the presented sound, and explicitly instructed to ignore the colors. Subjects were encouraged to minimize eye movement by maintaining central fixation to maximize hit chances and reduce noise; eye movement was free in the task nevertheless. After the AV presentation, the two symbols presented in the trial were shown again on the screen and a click sound prompted the observer to respond by pressing a keyboard button selecting one option (**Figure 1D**). After a choice, a high- or low-pitched feedback tone then indicated whether the response was correct or incorrect. In this task, accuracy was explicitly emphasized to participants over response speed, and responses made before the click were not accepted. Symbol pairs, colors, and auditory AM rates were equally distributed over 120 trials. In each task, trial order was randomized, and the percent correct performance was calculated and shown as feedback at the end. This value was multiplied by the number of points awarded at the immediately preceding Training task, resulting in the amount of tokens earned; for instance, 2000 points gained from Tr1 could be turned into a maximum of 2 tokens at a 100% correct rate at Tt1. Participants were informed of the number of tokens accumulated so far at the end of each Testing task. Testing tasks Tt1 and Tt2 were identical except that, as an incentive to improve their performance, at the start of Tt2 subjects were offered the possibility to pick (rather than be given) from an unspecified set of award options if they over-performed Tt1. At the end of the session participants either selected, or were assigned, a chocolate bar among different styles.

### 2.6 Data preprocessing

Data analysis was implemented in MATLAB 2018b (Natick, United States). EEG recordings from pre-experiment (auditory-only) sessions were downsampled to 1024 Hz, and to 512 Hz from main sessions. Head and reference sensor datasets were common average-referenced, after which DC offset was removed per channel. Signals were bandpass-filtered between 1 and 40 Hz with a 10-order elliptic infinite impulse response filter of 60 dB attenuation and 0.5 dB passband ripple. Single channel data were rejected in a blind manner according to a channel variance-based criterion (***Junghöfer et al., 2000***) with confidence coefficient τχ*_p_*=4, and the same procedure was performed separately for the external reference channels. To denoise EEG data (see below) and to address variable oculomotor behavior (see Electrooculogram data), recordings from the reference sensors were kept separate. To reduce artifact contamination, EEG data were decomposed by independent component analysis using FastICA (***Hyvärinen, 1999***). The two independent components with maximal broadband power were automatically selected and projected out of the data. A time-shifted principal component analysis (***de Cheveigné & Simon, 2007***) was applied to discard environmental signals recorded on the external reference sensors (±4 ms shifts). A sensor noise suppression algorithm (***de Cheveigné & Simon, 2008***) was applied in order to attenuate artifacts specific to any single channel (63 neighbors). EEG and electrooculogram (EOG) data were epoched. After epoching EEG data, the variance-based rejection procedure was again applied across single EEG channels and epochs’ timeseries per participant, resulting in less than 2.9% rejected channel-epoch timeseries form main sessions on average (subject range 2.7%–3.2%).

### 2.7 Data and statistical analysis

#### 2.7.1 Behavioral data

In Tr1 and Tr2 blocks, hit rates and response times were obtained per participant. In Tt1 and Tt2 blocks, d-prime (*d’*) scores indexing sensitivity to the audiovisual discrimination task were computed as the *z*-transform of the hit rates minus the *z*-transform of false positives to 9 Hz auditory stimulus presentations. *z*-scores were obtained by the normal cumulative distribution inverse function approximation, and *d’* values were obtained per condition, per subject. To ensure convergence, all hit rates and false alarm scores were approximated by the log-linear approach (***Stanislaw & Todorov, 1999***).

#### 2.7.2 Statistical analysis of behavioral data

All statistical analyses were performed with MATLAB R2018a, both with built-in functions as well as custom-written code. Effects were statistically assessed for confirmatory testing using linear mixed-effect (LME) models with the MATLAB function `fitlme`. The omnibus LME analyses evaluated the effects of independent ordinal variable “Stage” (Testing task 1 and Testing task 2), numeric variable “Reward” level (High, +80 points average; Low, +20 points; and No Reward, +0 points), and numeric variable “Uncertainty” (Probabilistic, 7 points standard deviation; Fixed, 0 points s.d.) on the dependent numeric variable (I: hit rate at Training; II: reaction time at Training; III: *d’* sensitivity at Testing). For Training tasks, the relevant “Flanker” variable (congruent, neutral, incongruent) was also included.

For Training data (*N*=32), the full model includes 16 fixed effects (Stage, Reward, Uncertainty, Flanker, and all higher-order interactions). To account for repeated measures within participants, the model includes random effects (intercept and slope) for each participant. In Wilkinson notation, the full general model is expressed as: *y* ∼ Stage × Reward × Uncertainty × Flanker + (Stage × Reward × Uncertainty × Flanker | participant).

For Testing data (*N*=31), the full model includes 8 fixed effects (same as above except Flanker, four interactions (Stage:Reward, Stage:Uncertainty, Reward:Uncertainty, Stage:Reward:Uncertainty) plus a nuisance fixed effect of participant handedness, to exclude the potential confounds of cerebral dominance asymmetries. Similarly to Training data, the model includes random effects (intercept and slope) for each participant in the between-subjects design involving two task difficulty levels as nuisance variable. The full general model is expressed as: *y* ∼ Stage × Reward × Uncertainty + Handedness + (Stage × Reward × Uncertainty | participant/difficulty).

Where a significant interaction was found, post hoc tests were carried out on with a Bonferroni-Holm correction to account for multiple comparisons. For reward learning effects observed at Tr2 or Tt2, subject-level association strength analyses were additionally performed. These analyses served to examine and illustrate the individual dependence of observers’ behavior with reward level as predictor. Beta (slope) parameters obtained by linear regression on individual subject data were then submitted to Student’s *t*-testing to evaluate the hypothesis that reward associations are established, i.e., association strength is significantly different from zero.

#### 2.7.3 Electrooculogram data

As variable saccades and gaze dwell may impact phase locking of the visual system to the image display elements, oculomotor behavior was evaluated via horizontal and vertical electrooculogram (EOG) recordings in the experimental sessions. For every trial in a Testing block, we computed the proportion of trial time when instantaneous recordings in any of the 4 EOG channels array exceeded a threshold. The threshold was determined per channel based on all recorded values across trials in the block. This proportion of excess activity was used as a proxy of the trial’s relative stability of sustained ocular fixation. To formally determine whether this oculomotor index had a confounding role on Testing stage findings about neural PLV, we (i) submitted it to the same statistical model, and (b) regressed it against neural PLV to evaluate significance of the predictor relationship.

#### 2.7.4 Audiovisual locking data analysis

We probe whether reward learning impacts the stability of locking responses to audiovisual temporal at the single trial level, under both probabilistic and fixed modes of learning. EEG channel data from Tt1 and Tt2 blocks were epoched by 6 s AV presentation segments, sub-epoched into 1 s partitions, then transformed into the frequency domain via a Fast Fourier Transform, resulting in six sub-epochs spectra per trial with 1 Hz resolution. Phase coherence analysis (***Nozaradan et al., 2012***; ***van Diepen & Mazaheri, 2018***) was used to measure temporal and spectral phase synchronization in EEG signals at the single trial level, via circular averaging of the corresponding phase angle timeseries across sub-epochs spectra. Resulting intra-trial phase locking values (PLV) were used to evaluate the stability of steady state responses in each trial as they become modulated by temporal lapses of attention (***Keitel et al., 2019***). PLVs were obtained by estimating peak sizes of phase coherence spectra at the target auditory stimulation rate plus the first and second harmonics (i.e., 9, 18, and 27 Hz in 9 Hz trials; alternatively, 11, 22, and 33 Hz). Harmonics were included (**Figure 2A**) as they may contain non-redundant neural representations that complement those directly linked to the base rate (***Jenkins et al., 2011***; ***Keitel & Müller, 2016***). Each PLV peak size was expressed as the ratio of the coherence at the target frequency bin divided over the average coherence at the two immediate neighbor bins. The PLV for a trial was defined as the average of the three sound-related peak frequency ratios across nine auditory-dominant (Fz, FC1, FCz, FC2, C3, C1, Cz, C2, C4, see **Figure 2B**), or when indicated visual-dominant (PO7, PO3, O1, POz, Oz, Iz, PO4, O2, PO8), EEG channels and expressed in decibels.

**Figure 2.**
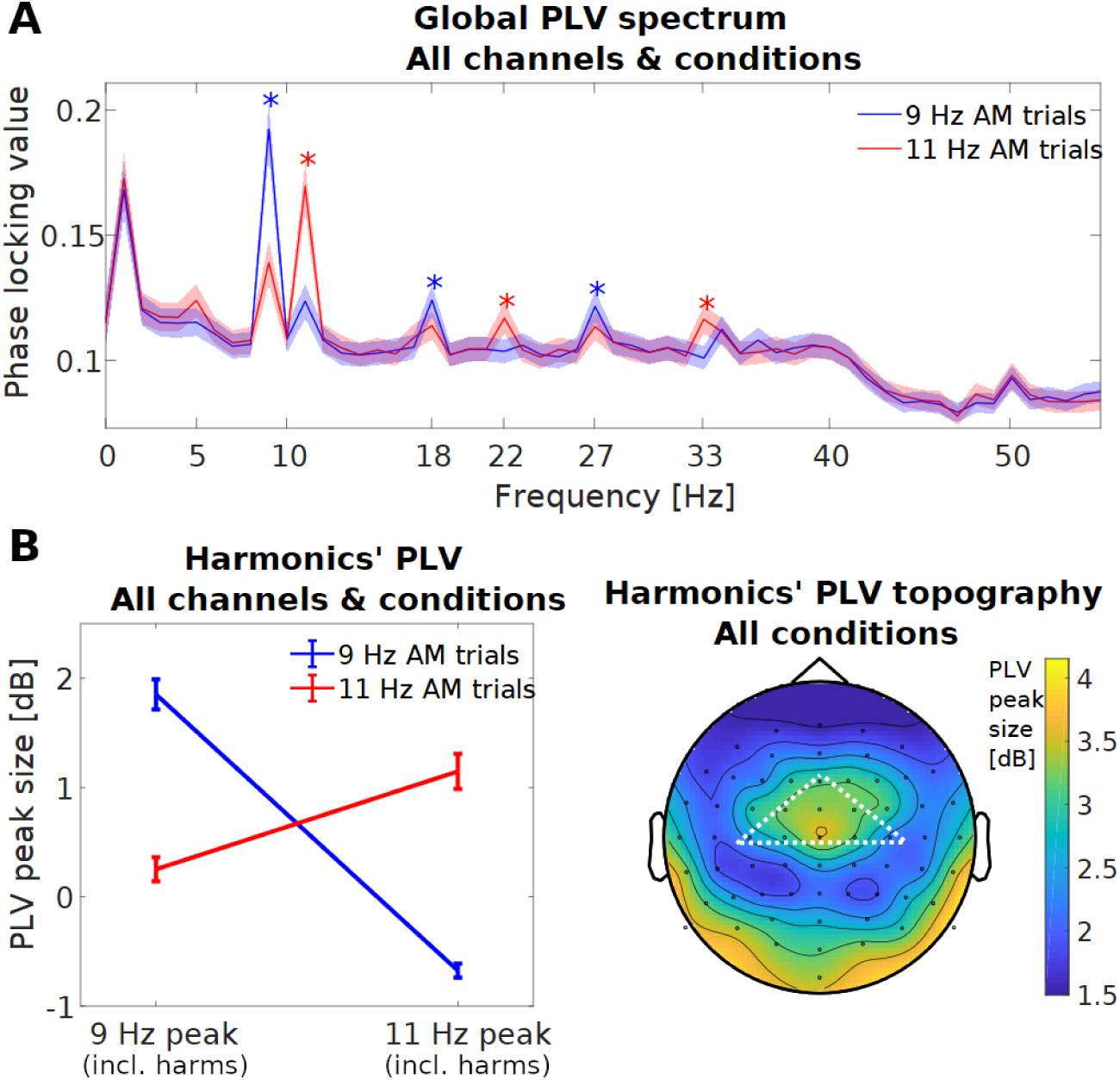
Single-trial estimation of temporal locking precision during AV scene presentations. (A) Intra-trial phase coherence spectra averaged across trials, conditions and channels, by sound modulation rate. Audiovisual (AV) presentations elicit characteristic frequency patterns in participants’ EEG responses, with significant phase-locking value (PLV) differences at the sound modulation rate and two harmonics (adjusted *p*-values ≤ 0.004). (B) Precision of temporal locking measured by the size of PLV peaks, estimated by referencing the crest phase coherence value relative to the immediate neighboring bin frequencies. Expressed in decibels, the average of these peak-to-baseline ratio estimates appears specific to the auditory stimulation rate (left). Error bars indicate ±1 standard error of the mean. Topographically, peak sizes show relatively precise temporal locking over the frontocentral scalp of the EEG array, in addition to occipital areas (right). The highlight indicates channels that were used to index AV locking to the auditory rate in subsequent analyses, unless otherwise indicated.

#### 2.7.5 Auditory locking data and statistical analysis

Finding of reward learning impacting precise temporal locking of audiovisual responses in auditory sites at Tt2 (see Results) prompted further investigation of the impact of learned reward effects on unimodal responses. In particular, we aimed to evaluate whether reward learning and performance effects similarly hold for auditory responses to the sound stream at this stage. Importantly, recordings of intra-trial coherent activity at the auditory channels include contributions to PLV from cross-modal activity locked to both visual flicker rates (**Figure 2A**). This is a consequence of visual and auditory streams maintaining a consistent phase lag within each trial, overall contributing to consistency across trial sub-epochs; AV lag is however variable between trials. Next, we evaluated phase consistency of auditory-locked activity devoid of visual contributions by using an inter-trial approach in combination with spatial source separation.

First, a spatial filter was constructed to emphasize reproducible auditory processing across subjects using an implementation of joint decorrelation (JD, ***de Cheveigné and Parra, 2014***; ***de Cheveigné and Simon, 2008***). The spatial filter weights were estimated simultaneously from all listeners’ recordings of the ‘auditory-only’ pre-experiment session, each organized in two 54 s epochs corresponding to 9 and 11 Hz presentations. Data were submitted to a first round of JD using as bias data the average across listeners in the two separate presentations. From the resulting spatial decomposition, the subspace resulting with the highest stimulus-evoked activity ratio across the two stimuli was extracted. This reproducible subspace, composed of the top 3 components, was projected back into sensor data, treating less reproducible activity across listeners as noise. Next, denoised unisensory data were filtered using an order-2 Butterworth filter in the 8–10 (for 9 Hz stimuli) or 10–12 Hz (11 Hz stimuli) bandpass region and averaged across participants. Denoised sensor data were submitted to a second round of JD using as bias data the filtered average across listeners in the two separate presentations. The most repeatable (top) component was consistent with auditory source activity and used together with the first step to decompose. The procedure provided two transformations: one to remove the spatial components least likely to reproduce across listeners, and another to optimize the signal to noise ratio of isolated stimulus-evoked narrowband activity (***de Cheveigné and Parra, 2014***) from the auditory system.

Second, the above transformations obtained from listening-only data were then applied to AV data. Specifically, 1) the first JD matrix was applied to single-trial Tt1 and Tt2 data; 2) the top 3 components were kept and projected back into sensor space; 3) the second JD matrix was applied to the denoised data; and 4) the top component was used for auditory tracking analyses. As only linear weightings of sensor data are applied, narrow-band spectral filtering of the main experiment epochs was not imposed. After obtaining auditory source estimates, inter-trial phase coherence (ITPC) measures were applied across trials. ITPC measures are sensitive to sample size difference biases, and were thus computed for each subject at 20 trials per reward level condition and subject. Because visual flicker phase lags varied from trial to trial relative to sound AM phase, the procedure effectively discarded visual oscillatory contributions to phase variability. In this case, only the first harmonic of each fundamental rate resulted in an obvious peak across listeners and conditions, phase variability analyses of auditory locking therefore did not include the second harmonic. For each reward level and subject, ITPC spectrum data (*N*=30) were collected at the sound presentation AM rate and its first harmonic, across both AM conditions in the Tt2 stage. ITPC data were regressed against reward level and resulting beta parameters were submitted to Student’s *t*-testing.

## 3. Results

We examined participants on a two-alternative forced choice (2AFC) AV coherence task before and after a reward conditioning visual procedure (**Figure 1**). The multistage process was used to determine whether the temporal precision tracking AV modulations during the 2AFC task (**Figure 2**) may be influenced by reward associations established through conditioning.

### 3.1 Reward manipulation incentivizes observer behavior during conditioning

To first determine whether the reward association was established, we inspected subject behavior before and during conditioning. Both training stages, consisting of a standard flanker’s task with color cues (**Figure 1**), resulted in performance that was assessed by linear mixed effects modeling. Hit rates and, separately, mean reaction times analyzed by the factors ‘Flanker’ (standard congruent, neutral, incongruent conditions), ‘Stage’, ‘Reward’, ‘Uncertainty’, and their interactions, first showed an effect of ‘Flanker’ condition in both hit rate (*F*_2,936_ = 9.83; *p* < 0.001) and reaction time data (*F*_2,996_ = 67.6; *p* < 0.001), as is expected for this task. While hit rates were not further significantly modulated by any other factor or interaction (*p* > 0.09; Supplementary Table 1), mean reaction times showed a significant main effect of ‘Stage’ (*F*_1,996_ = 11.5; *p* < 0.001). This main effect was qualified by (*i*) a significant interaction with ‘Flanker’ (*F*_2,996_=4.59; *p* = 0.010); (*ii*) a significant interaction with ‘Reward’ (*F*_1,996_ = 5.08; *p* = 0.024; Supplementary Table 2). Addressing the significant ‘Stage’ by ‘Flanker’ interaction, post-hoc analyses revealed that responses to neutral and incongruent flanker trials were faster in the second training stage (during conditioning, Tr2) compared to the first (before, Tr1; Supplementary Table 3). Post-hoc analyses conducted to address the significant ‘Stage’ by ‘Reward’ interaction additionally revealed that responses to Low (20 point average reward) and High reward trials (80 point average) were faster in Tr2 compared to Tr1 (Supplementary Table 4). Mean reaction times were not significantly modulated by any other factor or interaction in the omnibus test (*p* > 0.42).

The results suggest that participants speed up inhibitory responses as a function of learning the color payoff context, relevant to increasing task gains (**Figure 3A**), indistinctly of varying and fixed conditioning. Therefore, we expect that if reward-driven learning has been substantiated at conditioning (Tr2), then reaction times at that point may be expected to vary as a function of reward. For each participant, mean reaction time was regressed against expected reward indicated by color, evaluated at 0, 20 or 80 points, and individual association slope parameters were obtained. Faster responses to higher expected reward predict negative slopes, and a one-sided *t*-test confirmed that at this reward learning stage the average of participants’ slopes was significantly lower than zero (*t*(31) = −2.574, *p* = 0.008).

**Figure 3.**
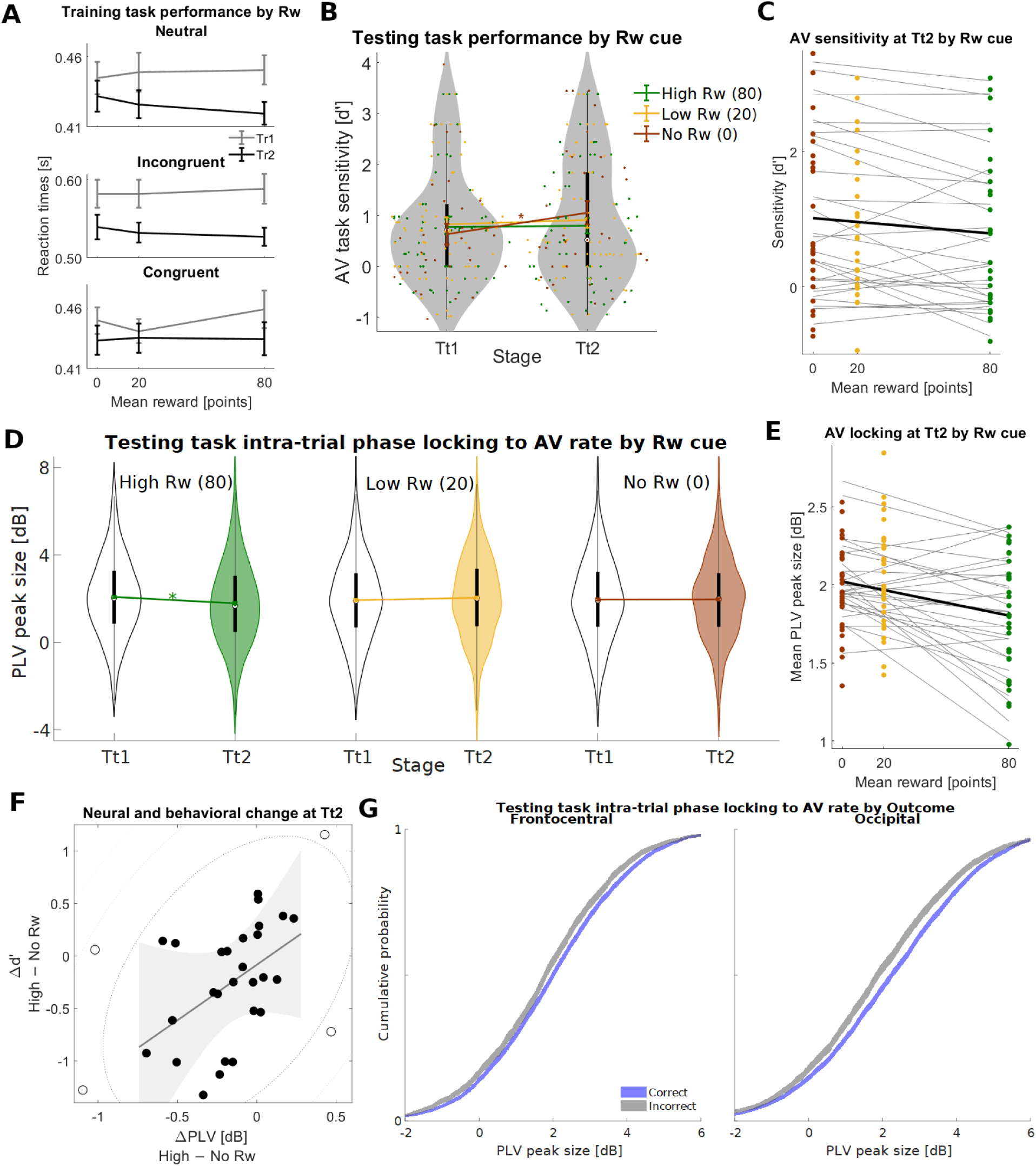
Behaviorally-relevant precise of locking responses to AV stimuli have stability interfered by reward-driven visual distraction. **(A)** In a standard flankers task subjects speed their inhibitory responses as a function of training and learned color-reward mappings, indicating consolidation of reward conditioning at the Tr2 stage. **(B)** Before and after conditioning subjects perform an independent two-alternative forced choice audiovisual (AV) correspondence task. Improved sensitivity performance at the latter Tt2 stage is contingent on lack of reward-associated color distractors. Dots denote subjects per condition (boxplot centroid: median; thick line: inter-quartile range). **(C)** Sensitivity data regressed against reward level shows decreased sensitivity at Tt2 to AV correspondence. **(D)** Neural results before and after conditioning show decreased intra-trial phase-locking to the common AV temporal modulation rate in the presence of high-reward-associated distractors, even as the latter’s temporal flicker is synced with visual targets. **(E)** Phase locking data regressed against reward level shows the retreat in stable temporal encoding as a function of prior reward context cued by distractors at Tt2. **(F)** The loss in temporal precision predicts subjects’ weaker behavioral sensitivity in the AV task (reward-driven distraction). Contour indicates distance from bivariate mean; data with *D_s_* ≥ 6 squared z-units are defined as outliers (open circles). Shaded region: 95% confidence intervals (CI) of linear regression model prediction. **(G)** 95% CI of the empirical cumulative distribution function of single-trial phase locking values show stable, precise locking as a relevant decision variable in the AV task. All single-trial data combined show that, across all conditions and stages, decreased stability of responses locked to the common temporal AV rate predict incorrect outcomes, measured at the frontocentral locations (left) but also at occipital sensors (right).

### 3.2 Cue-dependent sensitivity consistent with reward-driven distraction in AV task

Given an effective color-reward association, behavioral measures of AV discrimination sensitivity performance (*d’*) were next evaluated at the pre- and post-conditioning stages of AV testing (Tt1 & Tt2, respectively). Rather than aligning with task goals (as in the training tasks), conditioned peripheral color cues here are likely to interfere and lead observers to suboptimal performance. Participants observed the single-auditory/dual-visual streams conforming the target AV display before committing to a single visual option that matched the sound. Hence, performance measurements at Testing were not based on reaction time and were limited to sensitivity. Linear mixed effects analysis of *d’* by the relevant factors ‘Stage’, ‘Reward’ associated with the peripheral dots, ‘Uncertainty’ associated with the same, and their interactions, first confirmed a significant main effect of ‘Stage’ on AV sensitivity (*F*_1,301_ = 7.50; *p* = 0.007), which increased from Tt1 to Tt2 (**Figure 3B**) consistent again with practice effects. This main effect was qualified by a significant interaction with ‘Reward’ (*F*_1,301_ = 4.52; *p* = 0.034). *d’* values were not significantly affected further by any other factor or interaction (*p* > 0.19; Supplementary Table 5). To address the significant interaction, post-hoc analyses were conducted to evaluate the effect of stage across each reward condition, revealing a significant increase in sensitivity in Tt2 compared to Tt1 that was limited to No reward trials (*F*_1,31_ = 9.94; adj. *p* = 0.011; **Figure 3B**); for Low and High reward trials there was no significant change (*p* > 0.9; Supplementary Table 6). To clarify whether these findings may be interpreted as reward cues interfering with optimal performance, Tt2 sensitivity data were evaluated as a function of expected reward magnitude. For each participant, mean *d’* was regressed against expected reward signaled through color distractors, and individual association slope parameters were obtained (**Figure 3C**). Weaker sensitivity under higher expected reward associations at this testing stage also predict negative slopes, and one-sided *t*-testing confirmed that the mean of participants’ slopes was significantly below zero (*t*(30) = −2.324, *p* = 0.013).

The results thus are consistent with implicit reward learning transferred over the testing stage, and overall suggest that practice benefits at the second round may be offset or not available in the presence of rewarded-associated signals. To facilitate comparison with reward-driven distraction studies (***Rusz et al., 2020***) we additionally computed the standardized mean change effect size of d’ sensitivity measures at Tt2, contrasted between the High reward (mean *d’* = 0.80) and No Reward (mean *d’* = 1.05) conditions. Expressed in standard deviations, the parameter quantified how much of potential performance appears hindered by the reward cue. The estimated effect size of 0.192 result is moderate for visual reports but in line with cross-modal studies (***Rusz et al., 2020***).

### 3.3 Contextual reward cues disengage stable tracking of AV objects

Next we studied how induction of reward associations changed the intra-trial stability (temporal precision) of locking responses to key modulations of the AV scenes. Scalp-wide EEG activity during these presentations responded as expected to the stimulation design (**Figure 2**). To assess the temporal precision of locking responses, we estimated the peak-to-baseline ratios (peak sizes) of EEG channels’ PLV spectra at frequencies determined by auditory/target modulation rates (9, 18, and 27 Hz; alternatively, 11, 22, and 33 Hz). These estimates were found to map topographically to a distribution predominantly consistent with auditory activity (***Nozaradan et al., 2012***) and including occipital locking.

To study cross-modal VDAC effects on the temporal precision of tracking responses, PLV peak size data from frontocentral auditory channels were averaged (**Figure 2B**) and analyzed, on a trial-by-trial basis, by a full linear mixed effects model as in behavioral sensitivity analyses. We addressed oculomotor behavior as confounding variable, and submitted single-trial EOG data estimates of excess activity to the same analysis. While EOG data showed most trials had 10% or less presentation time involved with excess activity (Supplementary Figure 2), there was no significant effect or interaction on this variable ( *p* > 0.10; Supplementary Table 7). Furthermore, the distribution of EOG data did not predict PLV peak size estimates from AV responses (Supplementary Figure 2).

Analysis of PLVs by the factors of interest ‘Stage’, ‘Reward’, ‘Uncertainty’, and their interactions confirmed a significant interaction between ‘Stage’ and ‘Reward’ (*F*_1,7430_ = 4.29; *p* = 0.038) in these neural data. No other significant interaction or main effect was found (Supplementary Table 8). The significant interaction was analyzed in post hoc tests, revealing a significant decrease in PLV by ‘Stage’ at the High reward condition (*F*_1,2478_ = 13.63; adj. *p* = 6.8×10^−4^), with no evidence of change at the other reward conditions (Supplementary Table 9). The results suggest that when visual distractors cue previous reward to observers in a sustained fashion, the ability of the brain to precisely sustain temporal representations of target AV dynamics becomes suboptimal (**Figure 3D**).

As with behavioral data, we addressed whether the neural changes that were transferred via conditioning to the AV task may reflect reward magnitude. Single-trial Tt2 phase locking data were investigated as a function of expected reward signaled by color distractors, and individual association slope parameters were obtained (**Figure 3E**). Weaker phase locking by increasing expected reward associations similarly predict negative slopes in this case. One-sided *t*-testing confirmed that the average of participants’ slopes was significantly below zero (*t*(30) = −3.763, *p* = 3.7×10^−4^). The results indicate that visual distractors decreased the temporal precision of phase locked AV responses at frontocentral scalp, via their past associations with reward level.

### 3.4 Cue-dependent loss of stable tracking indexes observers’ reward-driven distraction

The finding of reward association effects at both behavioral and neural levels motivated the hypothesis that both may be related. We examined whether, at a subject level, the scale of changes to temporal encoding evidenced by PLV predicted the behavioral performance differences. Effect sizes (in *d’* and PLV) were defined by the separation between the High and No Reward conditions. At the Tt2 stage, participants’ PLV effects were significantly correlated with their behavioral differences (Spearman’s ρ = 0.473, [0.113 – 0.735 C.I.]; *p* = 0.007). Steeper losses of precise temporal locking in a participant predicted their greater loss of sensitivity in the task (**Figure 3F**). The correlation was robust to potential outliers (Shepherd’s Pi = 0.529; *p* = 0.010), and a robust regression analysis additionally confirmed its significance (*t*=2.48; d.f.=29; *p* = 0.019). By contrast, Tt1 stage (pre-conditioning) data failed to reveal a significant correlation between neural and behavioral effects (ρ = 0.228; [−0.171 – 0.556 C.I.]; *p*=0.22) in full, or after the removal of potential outliers (Pi = 0.159; *p* = 0.82).

### 3.5 Temporally stable selective tracking is a neural index of correct AV stream integration

The previous result suggests that visual reward-driven distraction may interfere with the tracking processes used to establish reliable temporal associations within the AV scene. VDAC deficits have been shown to reflect adverse sensory encoding or weak sensory evidence integration in at least sensory cortical regions (***Hickey & Peelen, 2015***; ***Pearson et al., 2022***). While our findings are consistent with spontaneous out of phase fluctuations within a trial that negatively impact perceptual decisions, they do not exclude the contribution by dedicated attention to the non-target flicker into less reliable decision-making. Such scenario may tie the loss of temporal precision to varying top-down selection under different reward cue contexts, rather than to reward-driven distraction directly, with selectivity being a primary predictor of outcome. To dissect the link of precise temporal locking to perceptual decisions, we examined whether this neural index determined trial outcomes in relation to reward learning. For this, we assessed whether reward learning, i.e, the statistical interaction between ‘Stage’ and ‘Reward’, varies according to the ‘Outcome’ of a trial (correct or incorrect) through an exploratory factorial design (see Materials and Methods). The test involved a LME model run on the same single-trial PLV data which included the trial factors ‘Stage’, ‘Reward’, and ‘Outcome’ of the result (correct versus incorrect response). If a trial’s successful outcome is supported by precise tracking of target modulations, a main effect of ‘Outcome’ may be expected. Temporal precision that is particularly affected in misses consistent with VDAC rather than misses generally, including those from attending nontargets, may be additionally shown through a triple interaction.

Confirming the expected interaction between ‘Stage’ and ‘Reward’ in the PLV data (*F*_1,7431_ = 10.15; *p* = 0.001), the analysis further revealed a significant main effect of ‘Outcome’ (*F*_1,7431_ = 6.69; *p* = 0.010). The effect was due to trials leading to correct responses showing greater PLVs than those leading to incorrect outcomes, across all experimental conditions, (**Figure 3G**). Nevertheless, no other significant interaction or main effect was observed (Supplementary Table 10). The findings indicate that down-modulations on PLV may predict incorrect trial outcomes generally, including but not limited those linked to VDAC. Through reward learning VDAC may effectively decrease PLV, regardless of whether this drop eventually leads to a correct or incorrect trial. The results therefore appear consistent with bottom-up interference forming temporally precise bimodal neural representations during the AV perceptual decision-making.

### 3.6 Auditory and supramodal activity is associated with loss of stable tracking elicited through VDAC

So far, stable locking to the target rate in frontocentral scalp within a trial may predict decision outcome. Because (*i*) locking changes by the visually cued VDAC may be present even in successful trials; and (*ii*) color distractors may not necessarily undermine visual temporal locking because they are strictly target-synced, we then evaluated activity from occipital regions. Such activity is also expected to reflect more directly the arbitration of the observer’s time-varying top-down selection across the dual flicker of central objects (***Kim et al., 2007***). Importantly, querying visual activity may inform us on the source of VDAC effects on temporal locking as the modality of origin conveying the reward-associated signals. Single-trial visual selection PLVs from occipital electrodes (see Methods), were submitted to the same statistical model as above. The analysis similarly showed a significant main effect of ‘Outcome’ (*F*_1,7431_ = 25.4; *p* = 4.7×10^−7^), indicating greater temporal locking to target rates in correct versus incorrect trials. This result confirms the impact that precise temporal representations of the target rate have on task performance across conditions and stages. Unlike frontocentral activity nevertheless, a significant interaction between ‘Stage’ and ‘Reward’ was not observed in these phase-locked responses (*F*_1,7431_ = 1.54; *p* = 0.21; Supplementary Table 11), or in steady-state response amplitudes (Supplementary Figure 3). The lack of evidence may appear consistent with observers’ unchanged visual locking when spatial attention deflects away from targets to peripheral reward-associated color distractors, which are in phase with the central target. Moreover, it raises the question that negative VDAC influences on reliability may appear within supra-modal locking or even at unimodal auditory locking.

To illustrate the spatial profile associated with the temporal locking effects associated with reward learning and outcomes, we performed single channel analyses of single-trial PLV data from which the spatial topographies of associated *F*-values could be derived. Using these maps, the analysis confirmed that the signal-to-noise ratio of ‘Outcome’ effects was optimal at the occipital areas (**Figure 4A**). This is consistent with the ability of participants to successfully localize and focus their visual responses, entrained themselves to the single target rate that. Remarkably, ‘Stage’ by ‘Reward’ interaction effects were shown to peak instead only within a limited subset of electrodes at frontocentral scalp locations (**Figure 4B**), consistent with in supramodal and/or auditory locking responses (**Figure 2B**). This result may suggest that sound-entrained, rather than flicker-entrained responses were primarily susceptible to modulation by VDAC.

**Figure 4.**
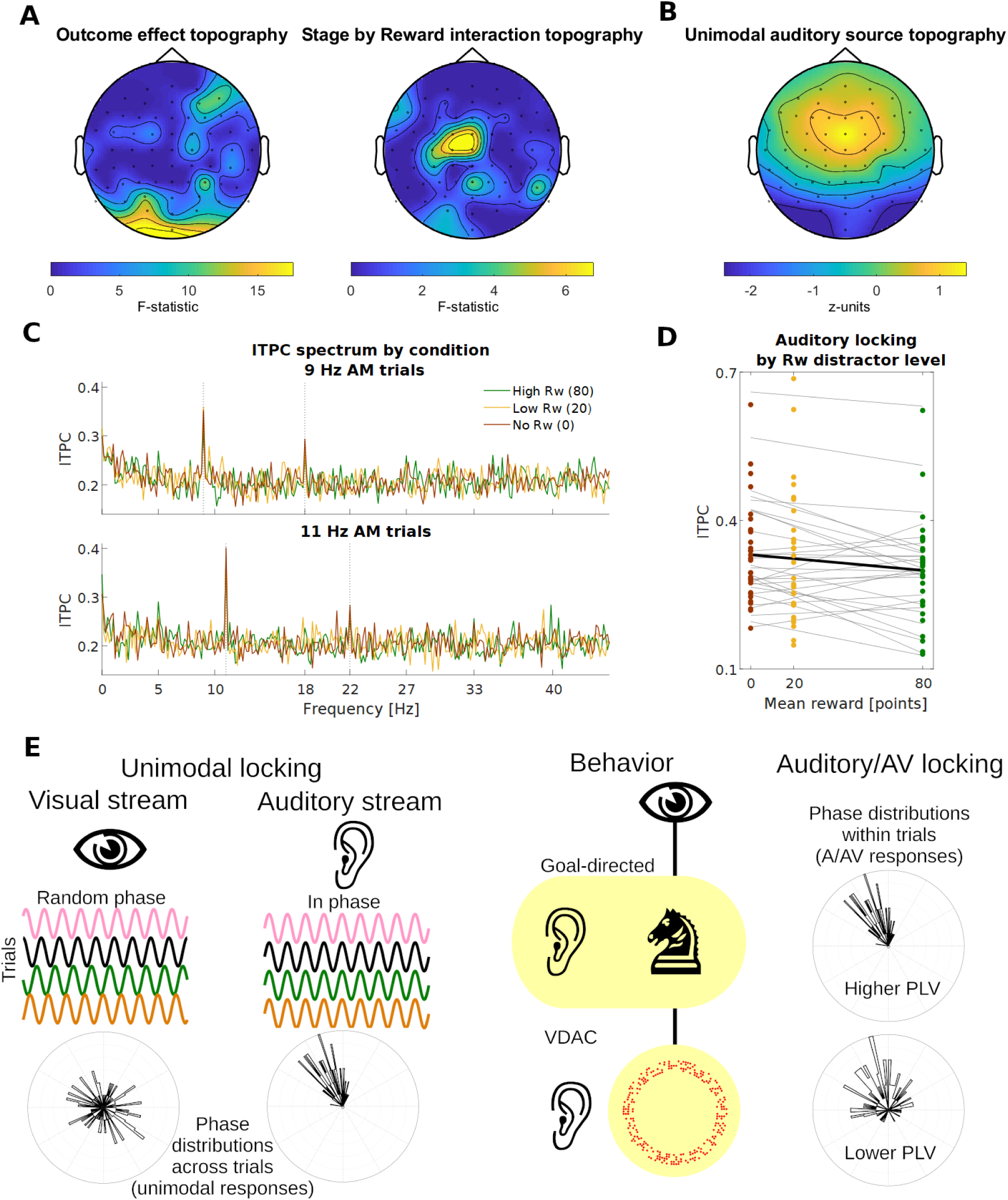
Reward-associated visual distractors decrease the temporal precision of auditory phase-locked activity. **(A)** Topography of repeated measures ANOVA *F*-values on single channel data, illustrating the localized strength of two relevant effects, reward learning and trial outcome. Largest effects of Outcome on single-trial sound/target rate phase-locking values are shown on occipital sensors (left), underscoring the relevance of selection of these rates for successful behavior. By contrast, the largest reward learning effect (Stage by Reward interaction) features a cluster of frontocentral electrodes (right), suggesting direct influence on auditory or supramodal phase locking. **(B)** Topography of the spatial component used for source separation of audiovisual (AV) data, trained from reproducible and narrowband activity that was obtained from additional auditory-only recordings of the same participants. **(C)** Unimodal auditory locking activity from Tt1 & Tt2 AV presentations, extracted via spatial source separation in combination with the inter-trial phase coherence method. Phase locking is clearly apparent at the sound amplitude modulation (AM) rate plus a harmonic (dashed lines) exclusively. **(D)** To evaluate the influence of reward-associated visual distractors, auditory inter-trial phase coherence values were regressed against reward level, demonstrating the decreasing effect by expected reward from color cues on the inter-trial temporal precision of sound tracking responses. **(E)** Conceptual illustration. *Left*: Target flicker stimuli match the AM in rate but have random phase. Because of this, associated unimodal visual responses will decohere when aligning responses by the acoustic phase. Noisy physiological responses, represented in phase diagrams, result in a relatively compact phase-locking distribution for unimodal auditory but which is absent in unimodal visual responses. *Right*: The observer’s behavior accommodates shifts of attentional sampling among the two candidate flickers in the dual visual stream. Visual attention to the correct flicker rate involves either the central object, in line with task requirements (top), or the periphery through reward-driven distraction (bottom). In the latter case, AV tracking supporting cross-modal integration decreases, suggesting intermodal competition (highlight).

### 3.7 Cue-dependent loss of temporal precision occurs within unimodal auditory tracking

In evaluating the hypothesis that auditory locking may lose temporal precision under VDAC, we acquired unimodal recordings of AM noise stimulation in the same listeners, which was used to train a series of spatial source separation filter (see Materials and Methods). Moreover, in each AV presentation during Tt1 and Tt2, the distractor flicker systematically varied in phase relative to the sound stream. Across-trial phase comparisons offer therefore an additional means to remove activity evoked by the dual visual flicker. Submitting unpartitioned (six-second) single-trial responses to source separation, in combination with inter-trial phase coherence (ITPC) estimation, first confirmed the absence of dual visual flicker responses (**Figure 4C**). Unimodal auditory locking, represented by peak ITPC values at the relevant AM stimulation frequencies (base plus first harmonic rates), was evaluated for the strength of reward associations by using linear regression. Here, loss of precise auditory locking under higher expected reward associations similarly predict negative slopes. Individual association slope parameters were obtained per participant (**Figure 4D**), and one-sided *t*-testing confirmed that the mean of participants’ slopes was significantly below zero (*t*(29) = −2.467, *p* = 0.010). The result is consistent with the notion that continuous cross-modal signaling of reward cue associations weaken the reliability with which temporal patterns are encoded in unconditioned unimodal cortex. Specifically here, visually carried reward signals operating as distractors directly decrease the quality of auditory entrainment to stimuli. This inter-modal suppressive effect appears to compromise the operation of low-level AV binding processes that are based on detecting common temporal dynamics.

## 4. Discussion

We interact with our surroundings more effectively when we integrate the flux of visual and auditory signals into veridically coherent audiovisual representations. Yet, the economy of attention remains earnest about obsolete reward associations (***Pearson et al., 2022***). This study asks whether these latter interfere with reliable AV processing, and demonstrates that conditioned visual signals may impact the temporal reliability of cortical responses to sound. When reward cues were being sampled, observers’ EEG responses showed reduced tracking precision to temporal modulations in the non-cued sense, as a function of learned reward. The change was followed by weaker task sensitivity detecting basic temporal correspondences in busy AV scenes.

Learned value associations modulate sensory cortical responses and alter the precision of neural representations of target input, whether from the same or different modalities as the value-associated distractor (***Itthipuripat et al., 2019***; ***Kim et al., 2021***; ***Pooresmaeili et al., 2014***; ***Antono et al., 2023***; ***Stănişor et al., 2013***). In the visual domain VDAC engages the reward system, the frontal eye field and posterior parietal cortical areas, where plasticity underpins changes in spatial priority (***Anderson, 2019***; ***Anderson et al., 2021***; ***Tankelevitch et al., 2020***; ***Bisley & Goldberg, 2010***; ***Markovic et al., 2014***; ***Kim et al., 2021***; ***Peck et al., 2009***). The parietal areas receive input encoding stimulus significance and mediate themselves shifts in the transmission of information according to behavioral relevance (***Sara, 2009***; ***Todd & Manaligod, 2018***; ***Mather et al., 2016***) and may feedback such input into sensory areas (***Antono et al., 2023***). Cross-modally, VDAC has been observed in temporal priming (***Anderson, 2016***), spatial co-localization (***Pooresmaeili et al., 2014***), and semantic coincidence paradigms (***Sanz et al., 2018***), and our present findings additionally implicate associations based on common temporal structure. Here, capture resulted in less stable audiovisual tracking activity consistent with systematic lapses of temporal sampling. Our results furthermore identified a weakened precision of unimodal auditory locking responses. Both findings suggest that recurrent lapses in attentional sampling originating from capture effects may create inter-modal competition which recurs over time.

For value distractors that are from a different modality than the target, bottom-up effects on unimodal processing may be observed relatively late (***Vakhrushev et al., 2023***; ***MacLean & Giesbrecht, 2015***; ***Tankelevitch et al., 2020***) and possibly no later than some visual-to-auditory influences operated through top-down attention (***Cervantes Constantino et al., 2023***). Delays possibly reflect fewer direct connectivity pathways between early sensory and frontal valuation areas. In intra-modal reward-driven capture, fMRI evidence points to mediation of the anterior intraparietal sulcus (IPS) communication to sensory area; in cross-modal conditions, communication between IPS and sensory areas may be instead relayed through the superior temporal sulcus (STS, ***Antono et al., 2023***). IPS and STS have a longstanding hypothesized role to establish together the spatiotemporal transformations required to form AV correspondences in the brain (***Miller and D’Esposito, 2005***). In addition, the IPS has been involved in audiovisual entrainment (***Bauer et al., 2020***, ***2021***; ***Lakatos et al., 2009***) and congruency based on normative spatial inference (***Rohe and Noppeney, 2015***). On the other hand, the temporal areas in or surrounding STS appear highly sensitive to AV temporal congruency, and appear to feedback synchrony effects automatically into early sensory areas (***Noesselt et al., 2007***; ***Lewis and Noppeney, 2010***; ***Atilgan et al., 2018***). A dichotomy of visually dominant, inference-led parietal versus auditory-dominant, timing correspondences-led temporal areas (***Werner and Noppeney, 2010***), could suggest that specialized temporal populations process AV time associations in a relatively automatic manner, while parietal populations may instead compute AV mappings based on spatial priority that is contingent on the observer’s behavior (***Noppeney, 2021***; ***Rohe and Noppeney, 2016***).

Distractors that capture attention away from targets in a stimulus-driven manner tend to increase task demands by interfering (***Tankelevitch et al., 2020***) while distractors that share the same target location typically boost performance (***Pooresmaeili et al., 2014***; ***Antono et al., 2023***; ***Grahek et al., 2021***; see reviewed in ***Rusz et al., 2020***). Findings of reduced rather than sustained locking appear consistent with suboptimal neural representations of sensory target input when salient reward cues are present (***Hickey & Peelen, 2015***). It is unclear how the neural representation of capturing distractors is enhanced because task resources are also dedicated to compute relative value (***Anderson, 2019***; ***Kim & Beck, 2020***; ***Liao & Anderson, 2020***; ***Anderson et al., 2021***). Such computation may as a result release from inhibition visual cortical regions encoding salient distractor representations to the effect that spatial and/or object-based target selectivities decrease (***Anderson et al., 2021***; ***Wang et al., 2015***; ***Hickey & Peelen, 2015***; ***Stănişor et al., 2013***). In addition, salience gains by distractors modulates the spectral and temporal profile of their corresponding neural activity. Cortical representations of visual stimuli exhibit more gamma-band power when stimuli appear more physically salient (***Fries, 2015***). Salience gains from neural activity in the temporal lobe also correspond with increased likelikood of AV binding at the perceptual level (***Mishra et al., 2007***; ***Balz et al., 2016***). Furthermore, reward conditioning boosts perceptual salience of unattended distractors (***Qin et al., 2021***) with temporal changes in neural responses a plausible means to reflect salience-based shifts in attentional sampling (***Jensen et al., 2012***). Taken together, these effects may imply that salience-led capture generates an adverse context of neural background activity for sustaining temporally precise target-following responses. Consistent with this, our findings suggest that visual reward distractor processing may limit timing acuity of audiovisual and auditory activity. We speculate this may be a manifestation of competition patterns from regions where unimodal visual dominant populations are interweaved with at auditory dominant and AV bimodal dominant populations, such as the temporal sulcus (***Beauchamp et al., 2004***). The balance between stimulus-following responses across these sub-populations, each modulated by corresponding salience-driven sub-threshold excitability dynamics, could then determine propagation of integrated or else segregated feedback into sensory networks (***Bauer et al., 2020***; ***Fries, 2015***; ***Keil & Senkowski, 2018***).

The present results furthermore suggest a continued ‘winner-take-all’ mode of capture (***Theeuwes, 2025***), as weaker reliability routes non-salient streams out of temporal correspondence and in effect segregated neural representations (**Figure 4E**). Robust alignment of neural activity with external rhythms may optimize perception as prediction of temporal and feature patterns facilitates targeted processing. Entrainment to rhythmic sensory input sets a phase relationship with one fluctuating but stable context to further sample, anticipate and interpret sensory content, while scaffolding the timecourse of activity across brain regions with diverse feature space mappings (***Lakatos et al., 2019***). When phases of neural oscillation excitability mismatch across different areas at similar times, sensory input may be segregated.

As in neural responses of predictable self-generated input (which are actively suppressed by the brain), background regularities could be exploited against them by the same ‘spotlight’ phase mechanism. Selectivity in entrainment models posits a combination of time- and feature-based phase resetting (***Lakatos et al., 2019***): enhanced representations of relevant objects follow from neural excitability in phase with most or all of its attended properties (timing plus other feature variables). By contrast, phase mismatch along any dimension promotes attenuation – whether mismatch in time correspondence (being out of synchrony) or, more critically, in mismatch in feature space (target and distractors are in synchrony but phase is matched to distractor features, ***Lakatos et al., 2019***).

The current study presents a number of methodological limitations. First, ensuring that target and distractors are in synchrony turned the latter strictly not task-irrelevant. Adopting distractors to perform is however a disadvantageous strategy because only central cues are reliable to determine a visual response. Importantly, syncing minimizes the likelihood that changes by any color distractors on visual temporal sampling are explicable by mismatched temporal phase. Findings that precise tracking from occipital areas is in general predictive of correct responses (**Figures 3G** and **4A**) suggest that oculomotor behavior relates to task goals without operating as a function of reward learning. For sensory sampling this setup minimizes distractor-led disruptions to visual time-locked responses. Second, saccade and gaze tracking recordings were not obtained, so their impact on visual time-locked responses is not determined. Key inference was therefore limited to auditory-locked activity, including frontocentral scalp responses (**Figures 3D** to **4B**) representing mixed signals from auditory, visual, and supramodal systems. Single-trial PLV data may be biased from variable gaze behavior if these contributions shift the target phase sets consistently across trial subepochs. Control analyses based on variability of the electrooculogram (an indirect measure) however suggest that eye activity was not significantly involved in single-trial PLV data. A third limitation stems from the fixed sequential design, which implies that causal effects of staging include order factors such as practice, fatigue, arousal, or time-on-task. Their causal separation from conditioning may require a control group following identical Tr1 and Tr2 tasks, for instance. Some of our findings suggest a dominance of conditioning over these factors, as for example absent modulatory effects on AV tracking by stage except at the high reward condition (**Figure 3D**). Such circumstances may be formally represented in the stage by reward interaction across statistical models, in addition to follow-up post hoc findings that are consistent with reward learning. Specifically, analyses of Tt2 stage data as a function of trained reward (e.g. **Figure 3E**) demonstrated sensitivity to the conditioned reward points task structure.

Despite these limitations, the break-up of robust AV binding in favor of value-driven inter-modal competition opens the question of the natural mechanisms by which incentive salience engages dynamic attentional sampling. Sustained top-down visuospatial attention is hypothesized to fluctuate rhythmically on regular sampling periods committed to a target, interleaved with disengagement that allows exploratory shifts (***Fiebelkorn & Kastner, 2019***; ***Lakatos et al., 2016***; ***Landau & Fries, 2012***; ***Todd & Manaligod, 2018***; ***Chen & Gong, 2022***). The latter periods represent temporary blind spots, referred to as ‘down-phases’ of the attentional cycle, harnessed by the visual system to explore parts of the scene for potential relevance. This framework may accommodate findings that reward cue signaling increases the likelihood of exploratory oculomotor behavior (***Rusz et al., 2020***), because recurrent exploitation of down phases may be warranted by salient distractors as their position changes. Securing these periods consistently may therefore provide a basic mechanism to sustain capture, because every returning cycle renews the ability to compete with goal-directed targets as they fluctuate across time.

## Author contributions

**Rodrigo Caramés Harcevnicow:** Formal analysis, Writing - Original Draft, Writing - Review & Editing, Visualization **Thaiz Sánchez-Costa**: Investigation, Data Curation, Writing - Review & Editing **Alejandra Carboni**: Software, Resources, Writing - Review & Editing, Supervision, Funding acquisition **Francisco Cervantes Constantino**: Conceptualization, Methodology, Software, Validation, Formal analysis, Investigation, Data Curation, Writing - Original Draft, Writing - Review & Editing, Visualization, Supervision, Project administration, Funding acquisition

## Data and code availability

Data and code supporting the findings of this study are openly available at the Open Science Framework at http://doi.org/10.17605/OSF.IO/SPJ9F.

## Acknowledgments

We thank Alex L. White and Germán A. Cipriani for code sharing, and Franco Caballero and Cecilia Biurrun for assistance with data collection.

