## Supplementary material for "Loss of precise auditory sampling as a sign of value-driven visual attentional capture"

### Supplementary Materials and Methods

#### Stimuli

**Video 1**. Sample AV testing (Tt1/Tt2) trial with a 9 Hz AM sound presentation. Both central figures shift locations every 1 second in this example. See Supplementary Fig. 1 for a video frame detail of the first two cycles of the target rate.

### Supplementary Figure

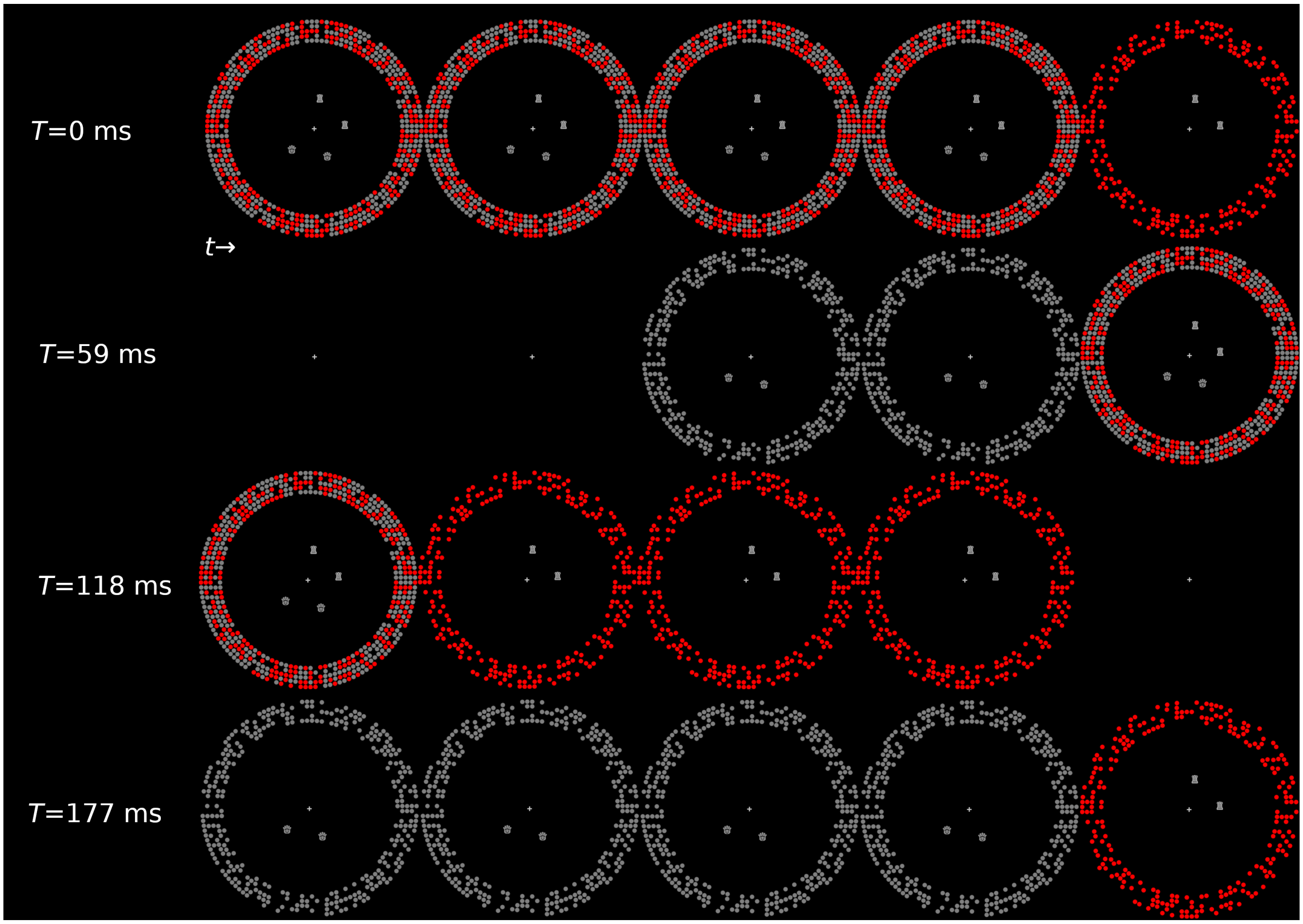


**Supplementary Figure 1.** Frame detail of the first 220 ms of Video 1. Time t step increases correspond to the 12 ms frame period approximately. Due to the frequency approximation approach (see Methods), the duty cycle of the 9 Hz set is 19 frames, and 15 frames for the 11 Hz set. The correct figure in every trial (rook in this example) syncs exclusively with the colored dot distractor set.

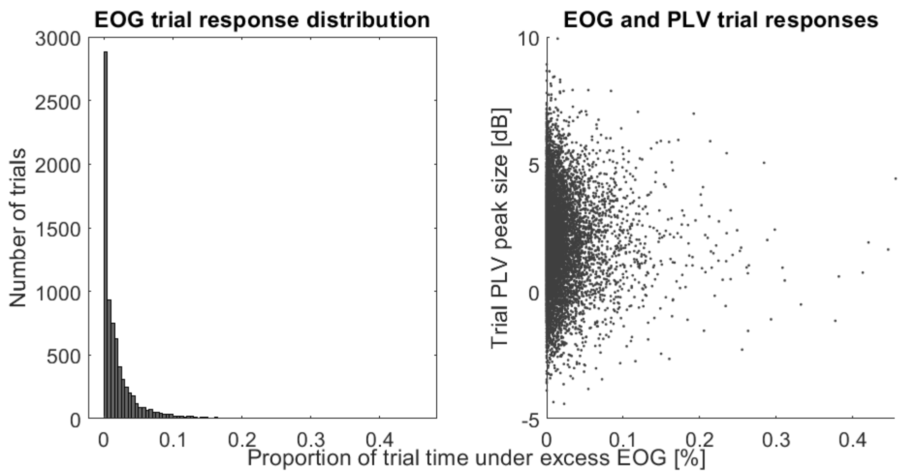


**Supplementary Figure 2.** Control analysis of eye-movement contributions to audiovisual (AV) phase-locking values (PLV). *Left*: Trial counts distribution as a function of total presentation time during which electrooculogram (EOG) electrodes yield excess activity (see Methods). Out of the 6 s presentation, most trials total 0.6 s (10%) or shorter periods of excess EOG activity. *Right*: Single trial AV PLVs as a function of excess EOG activity across conditions and participants. Evaluation of the within-subjects regression model PLV ~ EOG + (1|subject) indicates that excess EOG trial activity is not a significant predictor of AV trial PLVs ($\beta = -1.12$ ; $t(7437) = -1.72$ ; $p$=0.086).

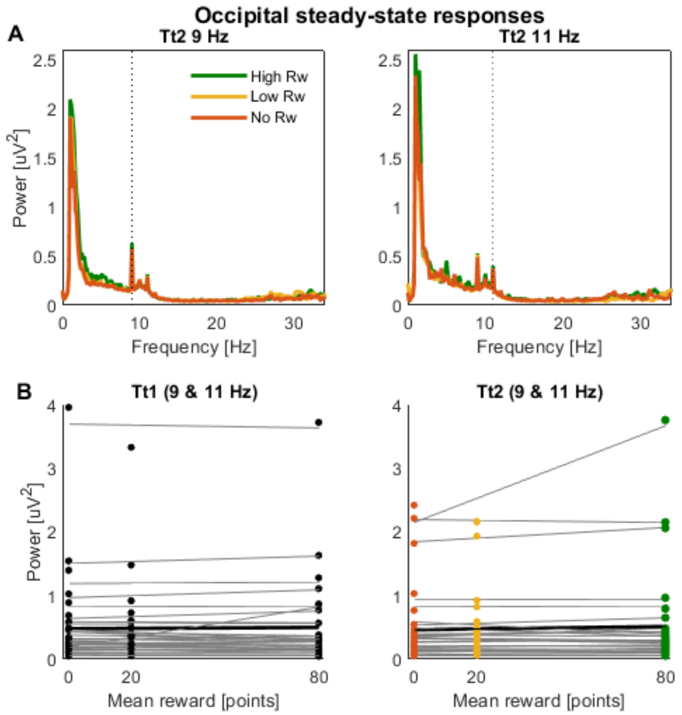


**Supplementary Figure 3.** Control analysis of occipital sensor steady-state responses at the target rate. (A) Grand-average power spectra of Tt2 responses by reward condition for trials with 9 Hz (left) and 11 Hz auditory presentation rates (right), consistent with evoked steady state responses to the dual flicker (AV) presentation, as a function of the auditory rate (dotted line). (B) Power amplitude across all trials by stage at the auditory rate by subject. Regressing steady-state responses against prior reward show that neither occipital activity from Tt1 (left) nor Tt2 (right) have regression slopes (grey: individual trendlines; black: average) significantly different from zero (Tt1: $t(31)=0.88$, $p=0.38$; Tt2: $t(31)=1.04$, $p=0.31$).

### Supplementary Results

#### Omnibus F-test analysis of behavioral performance (Training hit rates)

Model: HR ~ Flanker*Stage*Reward*Uncertainty + (Flanker*Stage*Reward*Uncertainty | participant)

| Term | F-value | *p*-value |
|---|---|---|
| Flanker | $F_{2,936}=9.83$ | **$6 \times 10^{-5}$** |
| Stage | $F_{1,936}=1.77$ | 0.18 |
| Reward | $F_{1,936}=2.28$ | 0.13 |
| Uncertainty | $F_{1,936}=2.81$ | 0.09 |
| Flanker × Stage | $F_{2,936}=1.12$ | 0.33 |
| Flanker × Reward | $F_{2,936}=1.33$ | 0.26 |
| Flanker × Uncertainty | $F_{2,936}=1.75$ | 0.17 |
| Stage × Reward | $F_{1,936}=0.65$ | 0.42 |
| Stage × Uncertainty | $F_{1,936}<0.01$ | 0.97 |
| Reward × Uncertainty | $F_{1,936}=0.60$ | 0.44 |
| Flanker × Stage × Reward | $F_{2,936}=0.93$ | 0.40 |
| Flanker × Stage × Uncertainty | $F_{2,936}=0.01$ | 0.99 |
| Flanker × Reward × Uncertainty | $F_{2,936}=0.33$ | 0.72 |
| Stage × Reward × Uncertainty | $F_{1,936}=0.85$ | 0.36 |
| Flanker × Stage × Reward × Uncertainty | $F_{2,936}=0.61$ | 0.55 |

Values in bold indicate *p*-values below 0.05 significance threshold

**Supplementary Table 1.** Omnibus analysis of subject hit rate behavioral data at Training stages 1 and 2 (flanker task).

**Omnibus F-test analysis of behavioral performance (Training reaction times)**

Model: RT ~ Flanker*Stage*Reward*Uncertainty + (Flanker*Stage*Reward*Uncertainty | participant)

| Term | F-value | *p*-value |
|---|---|---|
| Flanker | $F_{2,936}$=89.9 | **2x10$^{-36}$** |
| Stage | $F_{1,936}$=11.7 | **7x10$^{-4}$** |
| Reward | $F_{1,936}$=0.12 | 0.73 |
| Uncertainty | $F_{1,936}$=0.13 | 0.72 |
| Flanker × Stage | $F_{2,936}$=4.68 | **0.009** |
| Flanker × Reward | $F_{2,936}$=0.52 | 0.60 |
| Flanker × Uncertainty | $F_{2,936}$=0.36 | 0.70 |
| Stage × Reward | $F_{1,936}$=5.10 | **0.024** |
| Stage × Uncertainty | $F_{1,936}$=0.03 | 0.86 |
| Reward × Uncertainty | $F_{1,936}$=0.01 | 0.92 |
| Flanker × Stage × Reward | $F_{2,936}$=0.60 | 0.55 |
| Flanker × Stage × Uncertainty | $F_{2,936}$=0.21 | 0.81 |
| Flanker × Reward × Uncertainty | $F_{2,936}$=0.40 | 0.68 |
| Stage × Reward × Uncertainty | $F_{1,936}$=0.50 | 0.48 |
| Flanker × Stage × Reward × Uncertainty | $F_{2,936}$=0.87 | 0.42 |

**Supplementary Table 2.** Omnibus analysis of subject reaction time behavioral data at Training stages 1 and 2 (flanker task).

**Post-hoc analyses of training performance per stage and flanker type**

Model: $RT_j$ ~ Stage + (Stage | participant), where $j$ = {congruent, neutral, incongruent} flankers

| | Congruent flanker | | Neutral flanker | | Incongruent flanker | |
|---|---|---|---|---|---|---|
| | F-value | Adj. *p*-value | F-value | Adj. *p*-value | F-value | Adj. *p*-value |
| Stage | $F_{1,338}$=5.59 | 0.11 | $F_{1,338}$=10.6 | **0.007** | $F_{1,338}$=27.9 | **1.4x10$^{-6}$** |

P-values corrected for multiple comparisons through Bonferroni correction

**Supplementary Table 3.** Post hoc analyses of subject-level behavioral reaction time data by flanker and stage.

**Post-hoc analyses of training performance per stage and reward cue level**

Model: $RT_i$ ~ Stage + (Stage | participant), where $i$ = {+80, +20, +0} mean reward level

| | +80 (High reward) | | +20 (Low reward) | | +0 (No reward) | |
|---|---|---|---|---|---|---|
| | F-value | Adj. *p*-value | F-value | Adj. *p*-value | F-value | Adj. *p*-value |
| Stage | $F_{1,406}$=19.1 | **9.6x10$^{-5}$** | $F_{1,406}$=8.89 | **0.018** | $F_{1,202}$=4.62 | 0.20 |

P-values corrected for multiple comparisons through Bonferroni correction

**Supplementary Table 4.** Post hoc analyses of subject-level behavioral reaction time data by reward and stage.

### Omnibus F-test analysis of behavioral performance (Testing AV sensitivity)

Model: *d'* ~ Stage*Reward*Uncertainty + Handedness + (Stage*Reward*Uncertainty | participant\difficulty)

| Term | F-value | *p*-value |
|---|---|---|
| Stage | $F_{1,301}$=7.50 | **0.007** |
| Reward | $F_{1,301}$=1.23 | 0.27 |
| Uncertainty | $F_{1,301}$=0.09 | 0.77 |
| Stage × Reward | $F_{1,301}$=4.52 | **0.034** |
| Stage × Uncertainty | $F_{1,301}$=1.80 | 0.18 |
| Reward × Uncertainty | $F_{1,301}$=0.10 | 0.76 |
| Stage × Reward × Uncertainty | $F_{1,301}$=1.38 | 0.24 |
| Handedness | $F_{1,301}$=1.08 | 0.30 |

**Supplementary Table 5.** Omnibus analysis of subject sensitivity behavioral data at Testing stages 1 and 2 (AV task).

### Post-hoc analyses of testing performance per stage and reward cue level

Model: $d'_i$ ~ Stage + ( Stage | participant\difficulty), where $i$ = {+80, +20, +0} reward levels

| | +80 (High reward) | | +20 (Low reward) | | +0 (No reward) | |
|---|---|---|---|---|---|---|
| | F-value | Adj. *p*-value | F-value | Adj. *p*-value | F-value | Adj. *p*-value |
| Stage | $F_{1,122}$=0.03 | >0.99 | $F_{1,122}$=0.42 | >0.99 | $F_{1,60}$=9.94 | **0.008** |

P-values corrected for multiple comparisons through Bonferroni correction

**Supplementary Table 6.** Post hoc analyses of subject-level behavioral d-prime data by reward and stage.

**Omnibus F-test analysis of oculomotor behavior (Testing excess EOG activity)**

Model: EOG ~ Stage*Reward*Uncertainty + Handedness + (Stage*Reward*Uncertainty | participant\difficulty)

| Term | F-value | *p*-value |
|---|---|---|
| Stage | $F_{1,7430}$=0.42 | **0.52** |
| Reward | $F_{1,7430}$<0.01 | 0.99 |
| Uncertainty | $F_{1,7430}$=0.21 | 0.64 |
| Stage × Reward | $F_{1,7430}$=2.63 | **0.11** |
| Stage × Uncertainty | $F_{1,7430}$=0.07 | 0.79 |
| Reward × Uncertainty | $F_{1,7430}$=0.31 | 0.58 |
| Stage × Reward × Uncertainty | $F_{1,7430}$=1.06 | 0.30 |
| Handedness | $F_{1,7430}$=0.75 | 0.39 |

**Supplementary Table 7**. Omnibus analysis of subject EOG trial data at Testing stages 1 and 2 (AV task).

### Omnibus F-test analysis of single-trial locking activity (Testing AV phase locking values)

Model: PLV ~ Stage*Reward*Uncertainty + Handedness + ( Stage*Reward*Uncertainty | participant\difficulty)

| Term | F-value | *p*-value |
|---|---|---|
| Stage | $F_{1,7430}$=1.64 | 0.20 |
| Reward | $F_{1,7430}$=0.68 | 0.41 |
| Uncertainty | $F_{1,7430}$=0.19 | 0.67 |
| Stage × Reward | $F_{1,7430}$=4.29 | **0.038** |
| Stage × Uncertainty | $F_{1,7430}$=0.74 | 0.39 |
| Reward × Uncertainty | $F_{1,7430}$=0.32 | 0.57 |
| Stage × Reward × Uncertainty | $F_{1,7430}$=0.01 | 0.90 |
| Handedness | $F_{1,7430}$=0.96 | 0.33 |

**Supplementary Table 8.** Omnibus analysis of subject single-trial PLV peak size data at Testing stages 1 and 2 (AV task).

### Post-hoc analyses of training stage on single-trial AV PLV, per reward cue level

Model: $PLV_i$ ~ Stage + (Stage | participant\difficulty), where $i$ = {+80, +20, +0} reward levels

| | +80 (High reward) | | +20 (Low reward) | | +0 (No reward) | |
|---|---|---|---|---|---|---|
| | F-value | Adj. *p*-value | F-value | Adj. *p*-value | F-value | Adj. *p*-value |
| Stage | $F_{1,2478}$=13.63 | **$6.8x10^{-4}$** | $F_{1,2477}$=1.58 | 0.63 | $F_{1,2478}$<0.01 | >0.99 |

P-values corrected for multiple comparisons through Bonferroni correction

**Supplementary Table 9.** Post hoc analyses of single-trial auditory PLV data by reward and stage.

### Exploratory F-test analysis of single-trial locking activity (Testing AV phase locking values)

Model: PLV ~ Stage*Reward*Outcome + (Stage*Reward*Outcome | participant\difficulty)

| Term | F-value | *p*-value |
|---|---|---|
| Stage | $F_{1,7431}$=1.80 | 0.18 |
| Reward | $F_{1,7431}$=0.73 | 0.40 |
| Outcome | $F_{1,7431}$=6.69 | **0.010** |
| Stage × Reward | $F_{1,7431}$=10.15 | **0.001** |
| Stage × Outcome | $F_{1,7431}$=0.56 | 0.46 |
| Reward × Outcome | $F_{1,7431}$=0.28 | 0.60 |
| Stage × Reward × Outcome | $F_{1,7431}$=0.20 | 0.66 |

**Supplementary Table 10.** Exploratory re-analysis of auditory PLV peak size data, including outcome as predictor.

**Exploratory F-test analysis of single-trial locking activity (Testing AV phase locking values) at visual channels**

Model: PLV(occipital) ~ Stage*Reward*Outcome + (Stage*Reward*Outcome | participant\difficulty)

| Term | F-value | *p*-value |
|---|---|---|
| Stage | $F_{1,7431}$=3.49 | 0.06 |
| Reward | $F_{1,7431}$=1.82 | 0.18 |
| Outcome | $F_{1,7431}$=25.4 | **$4.7\times10^{-7}$** |
| Stage × Reward | $F_{1,7431}$=1.54 | 0.21 |
| Stage × Outcome | $F_{1,7431}$=0.07 | 0.79 |
| Reward × Outcome | $F_{1,7431}$=2.25 | 0.13 |
| Stage × Reward × Outcome | $F_{1,7431}$=0.02 | 0.90 |

**Supplementary Table 11.** Analysis of visual target PLV peak size data, including outcome as predictor.
